# Primary cilia promote resistance to EGFR tyrosine kinase inhibitor, osimertinib, in non–small cell lung cancer

**DOI:** 10.64898/2026.03.03.709408

**Authors:** Li Wang, Anuradha Pandit, Sk. Kayum Alam, Rhiannon Skauge, Sergio A. Gradilone, Luke H. Hoeppner

**Affiliations:** The Hormel Institute, University of Minnesota, Austin, MN, USA; Masonic Cancer Center, University of Minnesota, Minneapolis, MN, USA

## Abstract

Patients with advanced non–small cell lung cancer (NSCLC) and mutations in epidermal growth factor receptor (EGFR) benefit from EGFR tyrosine kinase inhibitors (TKIs). Osimertinib, a third-generation EGFR TKI, is standard first-line therapy for EGFR-mutated NSCLC, but most patients develop resistance to it. Here, we demonstrate that increased formation of primary cilia, microtubule-based sensory organelles, is associated with osimertinib-refractory NSCLC progression. EGFR-mutated, osimertinib-resistant human NSCLC cells had increased cilia formation and acetylation of α-tubulin and reduced histone deacetylase 6 (HDAC6) activity compared to their osimertinib-sensitive counterparts. HDAC6 inhibition increases cilia formation in osimertinib-sensitive NSCLC cells, and overexpression of exogenous HDAC6 sensitized osimertinib-resistant NSCLC cells to osimertinib’s anti-proliferative effects. Because intraflagellar transport (IFT) proteins are essential for primary cilium formation and function, we knocked down IFT88 in osimertinib-resistant NSCLC cells, which reversed osimertinib resistance in orthotopic and subcutaneous mouse models of lung cancer. Mechanistically, increased sodium influx during osimertinib-induced inhibition of EGFR signalling promotes cilia formation through sustained HDAC6 inactivity and greater α-tubulin acetylation. Inhibition of sodium influx with dibutyryl-cAMP decreased cilium formation, increased sensitivity to osimertinib, and reduced tumor progression in mice bearing osimertinib-resistant lung tumors. Collectively, our findings suggest that enhanced primary cilium formation mediates EGFR TKI resistance and that targeted inhibition of ciliogenesis may prevent or overcome osimertinib resistance.

---

Non–small cell lung cancer (NSCLC) is the leading cause of cancer-related death worldwide^1^. Oncogenic driver mutations in epidermal growth factor receptor (EGFR) that confer sensitivity to tyrosine kinase inhibitors (TKIs) are the most frequently occurring targetable alteration in NSCLC. Advanced NSCLC patients with mutations in EGFR benefit from EGFR TKIs; however, the development of resistance to NSCLC therapy is a major problem. Although EGFR mutation–positive patients respond well to first- and second-generation EGFR TKIs (gefitinib, erlotinib, afatinib), most patients develop resistance to these EGFR TKIs within 8-12 months and then experience rapid disease progression^2,3^. In ∼50% of these patients, a T790M secondary mutation occurs in EGFR^4,5^. Osimertinib, a third-generation EGFR TKI targeting the T790M mutation and primary activating EGFR mutations, was first approved for second-line treatment to overcome resistance to the first-and second-generation TKIs. However, only 61% of patients with T790M mutations respond to osimertinib, and in responding patients, cancer progression typically occurs again in <10 months^6^. In 2018, osimertinib was approved as a first-line treatment for EGFR-mutated lung cancer. First-line osimertinib exhibits superiority to other EGFR TKIs^7,8^, but most tumors inevitably acquire resistance to it, and patients are not cured. Patients with osimertinib-refractory lung adenocarcinoma have demonstrated amplification of MET in 15% of cases and C797S and L718Q EGFR mutations in 7% and 2% of cases, respectively^9,10^, but the mechanism of resistance is usually not known. Identifying therapeutic strategies to prevent or overcome resistance to first-line osimertinib remains an urgent unmet clinical need.

Primary cilia are microtubule-based sensory organelles that detect stimuli in the extracellular environment. The cilium projects from the apical surface of epithelial cells, and it is composed of a basal body anchored under the cell membrane and an attached axoneme that extends outside the cell surface^11^. The microtubules in primary cilia are arranged in a 9 + 0 configuration with nine outer microtubule doublets and no central microtubules, unlike motile cilia that consist of nine outer doublets and two single central microtubules in a 9 + 2 format^11^. Primary cilia house oncogenic molecules. These antenna-like organelles act as key signaling hubs of the Wnt, Hedgehog, MAPK/ERK, Notch, and receptor tyrosine kinase pathways^12,13^. As an important member of the receptor tyrosine kinase family, EGFR is involved in cell cycle control, migration and differentiation through the concentrated EGFR component on the axonemes of primary cilia^12,14^. EGFR suppresses ciliogenesis by directly phosphorylating the deubiquitinase USP8 on Tyr-717 and Tyr-810, which stabilizes the trichoplein-Aurora A pathway, thereby promoting cell proliferation^15^. In pathology, the relation of EGFR and ciliation has been widely studied in polycystic kidney disease. EGFR trafficking and cilia resorption is regulated by histone deacetylase 6 (HDAC6) through deacetylation of α-tubulin in cystic epithelial cells^16^. EGF also activates TRPP2/TRPV4, growth-associated amiloride-sensitive sodium-hydrogen exchangers, and ERK to increase cell proliferation and cystogenesis^17,18^.

Links between cilia structure/function and tumorigenesis have been reported^19^. Loss of cilia has been associated with initiation of some cancers, but increased ciliogenesis promotes oncogenesis and cell survival in other cancers^19^. Cilia may mediate drug resistance mechanisms in several cancer types^20^. NSCLC cells resistant to the first-generation EGFR TKI erlotinib exhibited more cilia than their erlotinib-sensitive counterparts^20^, which supports our rationale for investigating whether ciliogenesis is associated with osimertinib-refractory NSCLC progression.

Deciliation of cancer cells is promoted by overexpression of EGFR, based on EGFR-associated activation of KRAS and PI3K/AKT, or by activating mutations in KRAS, which is one of the potent mechanisms promoting cancer development. In contrast to their normal counterparts, cilia-deficient cholangiocytes show persistent EGFR activation because of impaired receptor degradation, a form of positive feedback^21^. However, for other types of cancer, like glioblastoma multiforme (GBM), the relationship between ciliation and tumor progression is ambiguous. One study established that ciliation is retained in GBM^22^, but another reported that ciliogenesis was disrupted at early stages of tumor development, with structural defects in the residual cilia^23^. Thus, the same type of cancer with distinct drivers shows different characteristics in ciliation. Medulloblastomas arising from constitutively active SMO, a Sonic Hedgehog (SHH) receptor, require cilia to promote their growth, while medulloblastomas arising from constitutively active GLI2, a molecule downstream of SHH signaling, requires loss of cilia to support its progression by eliminating signals inhibiting GLI2^24^. Thus, the role of cilia in promoting tumor progression or inhibiting tumor growth is disputed and likely context dependent. In addition to factors such as different tumor types and distinct genetic drivers of cancer, differing roles of cilia may also be caused by the interaction of cancer cells with multiple cell types in the tumor microenvironment, such as tumor-infiltrating immune cells, endothelial cells, and cancer-associated fibroblasts^25^.

Although the association between primary cilia and lung cancer has been investigated, it remains incompletely understood. An early study of the ChaGo human lung cancer cell line reported an absence of ciliation under basal conditions, although treatment with 2% DMSO induced ciliogenesis^26^. Another ciliation study on a total of 492 lung carcinomas used immunohistochemical analysis against a marker of primary cilia, ARL13B, to demonstrate that primary cilia are frequently present in SCLC but not in NSCLC. These primary cilia–positive cells showed activation of the Hedgehog signaling pathway^27^. A separate study demonstrated that cilia genes were upregulated in NSCLC^28^, and this finding was also verified by another study of human cancer samples^29^. These discrepancies may be caused by differing origins of cell lines, various stages of disease progression, and distinct lung cancer types. In mouse lung cancer models treated with cisplatin, a widely used chemotherapeutic agent, increased disruption of cilia was detected as ciliary proteins and fragments in the bronchoalveolar lavage fluid^30^. Human A549 cells that had acquired resistance to cisplatin exhibited increased ciliogenesis^20^. Similarly, increased primary cilia formation has been shown to promote the induction of acquired resistance to KRAS^G12C^ inhibitors^31^, and NSCLC cells resistant to 1^st^-generation EGFR TKI erlotinib exhibited greater cilia frequency than their erlotinib-sensitive counterparts^20^. Alterations in ciliation appear to be associated with changes in ciliary molecular composition, leading to enhanced activation of the Hedgehog signaling pathway^20,31^. Investigating whether cilia formation on NSCLC cells plays an important role in the development of osimertinib-refractory NSCLC progression represents an important, untested pursuit. The molecular basis of osimertinib resistance has yet to fully emerge because 1st-line osimertinib treatment became standard of care in the US relatively recently in 2018.

In this study, we report that the number of cilia is upregulated in NSCLC cells with acquired osimertinib resistance. This change is consistent with reduced HDAC6 activity and increased acetylation of α-tubulin. Inhibition of cilia through HDAC6 overexpression or intraflagellar transport 88 (IFT88) knockdown reversed osimertinib resistance in NSCLC cells. Consistent with these in vitro findings, genetic inhibition of cilia in orthotopic or subcutaneous lung cancer models indicated that reduced ciliogenesis increased the sensitivity of resistant NSCLC to osimertinib as marked by slower tumor growth. Further investigation showed that increased sodium influx during osimertinib-induced inhibition of EGFR signaling stimulates cilia formation through prolonged HDAC6 inactivation and enhanced α-tubulin acetylation. Pharmacologic suppression of sodium influx with dibutyryl-cAMP reduced cilium formation, restored sensitivity to osimertinib, and suppressed tumor progression in mouse models of osimertinib-resistant lung cancer. Taken together, our results suggest that targeted inhibition of ciliogenesis has the potential to prevent or overcome osimertinib resistance.

## Results

### Osimertinib-resistant NSCLC cells exhibit more primary cilia

To assess the role of ciliogenesis in the development of resistance to the EGFR TKI osimertinib in EGFR-mutated human NSCLC cells, we stained the cells for markers of primary cilia, ARL13B and ɣ-tubulin. We observed a greater number of primary cilia in human EGFR-mutated, osimertinib-resistant (OR) H1975-OR cells (i.e., H1975 R2 cells^32^) compared to their parental osimertinib-sensitive counterparts, H1975-OS (**Fig. 1a-b**). Given that increased acetylation of α-tubulin and reduced HDAC6 are associated with ciliogenesis^33–35^, we assessed these markers of primary cilia formation by immunoblotting. H1975-OR cells had decreased HDAC6 protein expression and more acetylation of α-tubulin relative to the H1975-OS cells (**Fig. 1c; Supplementary Fig. 1a-b**). To confirm these findings, we performed similar immunofluorescence and immunoblotting studies in another pair of human OS and OR NSCLC cells. HCC827-OR cells showed more cilia formation, more acetylation of α-tubulin, and less HDAC6 than HCC827-OS cells (**Fig. 1d-f; Supplementary Fig. 1c-d**). Collectively, our findings suggest that enhanced ciliogenesis is associated with osimertinib-refractory NSCLC progression.

**Figure 1.**
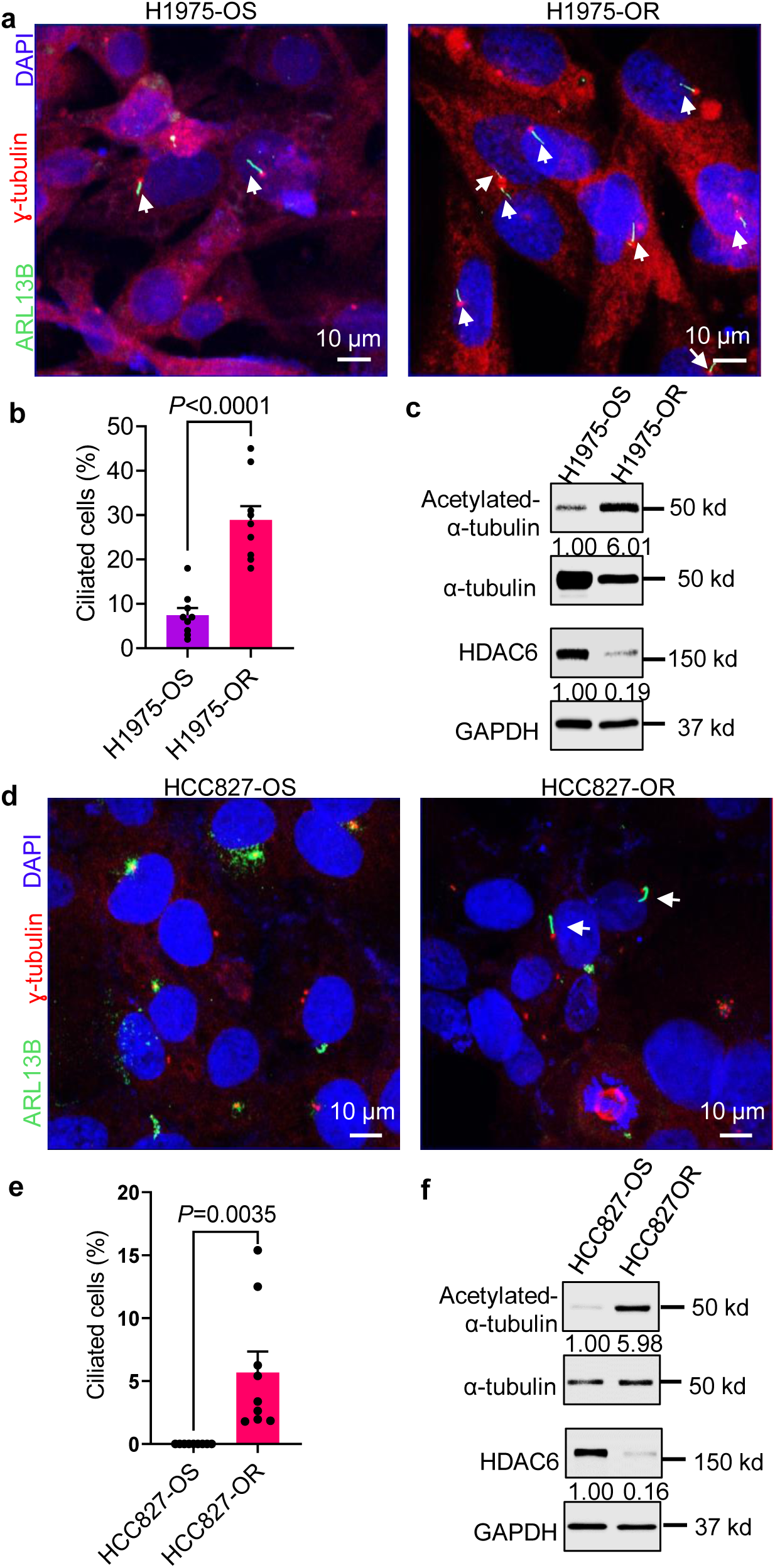
Increased primary cilium frequency in EGFR TKI osimertinib-resistant NSCLC cells. **a, b, d, e**, Serum-starved H1975 osimertinib-sensitive (H1975-OS) & H1975 osimertinib-resistant (H1975-OR) cells (**a, b**) or HCC827 osimertinib-sensitive (HCC827-OS) & HCC827 osimertinib-resistant (HCC827-OR) cells (**d, e**) were fixed; subjected to immunofluorescence staining to detect ARL13B (green) to mark cilia, ɣ-tubulin (red) for centrioles, and DAPI (blue) for the nuclei; and imaged on a Zeiss LSM900. **b,** Quantification of **a**. **e,** Quantification of **d**. Three biological replicates were performed, and representative data is shown. Unpaired t-test. **c, f,** Serum-starved osimertinib-sensitive (H1975-OS, HCC827-OS) and -resistant (H1975-OR, HCC827-OR) cells were lysed and immunoblotted with the indicated antibodies. Results of a densitometry analysis are shown beneath the bands. The depicted blot is representative of three experiments.

### HDAC6 regulates ciliogenesis in NSCLC cells

Given our observation that HDAC6 inactivation is associated with increased ciliogenesis in osimertinib-resistant NSCLC cells, we hypothesized that pharmacologic inhibition of HDAC6 in osimertinib-sensitive NSCLC cells would promote ciliogenesis. To test this hypothesis, we treated osimertinib-sensitive NSCLC cells with a potent and selective chemical inhibitor of HDAC6, ACY-1215 (i.e., ricolinostat)^36,37^, and assessed markers of ciliogenesis by immunofluorescence. H1975-OS and HCC827-OS cells both exhibited greater cilia frequency upon HDAC6 inhibition treatment than upon vehicle treatment (**Fig. 2a-b, d-e**). - Immunoblotting experiments revealed that pharmacologic inhibition of HDAC6 reduced HDAC6 protein expression and increased acetylation of α-tubulin in both H1975-OS and HCC827-OS cells (**Fig. 2c, f**). To determine whether overexpression of HDAC6 protein in osimertinib-resistant cells sensitizes them to the drug and reduces their primary cilia formation, we transiently overexpressed HDAC6 in H1975-OR cells (**Fig. 3a**). CellTiter-Glo® proliferation assays indicated that overexpression of HDAC6 sensitized them to osimertinib’s anti-proliferative effects (**Fig. 3b**). Overexpression of HDAC6 also inhibited ciliogenesis (**Fig. 3c-d; Supplementary Fig. 2**). Taken together, our findings suggest that inactivation of HDAC6 plays a key role in the increased ciliogenesis associated with osimertinib-refractory lung cancer progression.

**Figure 2.**
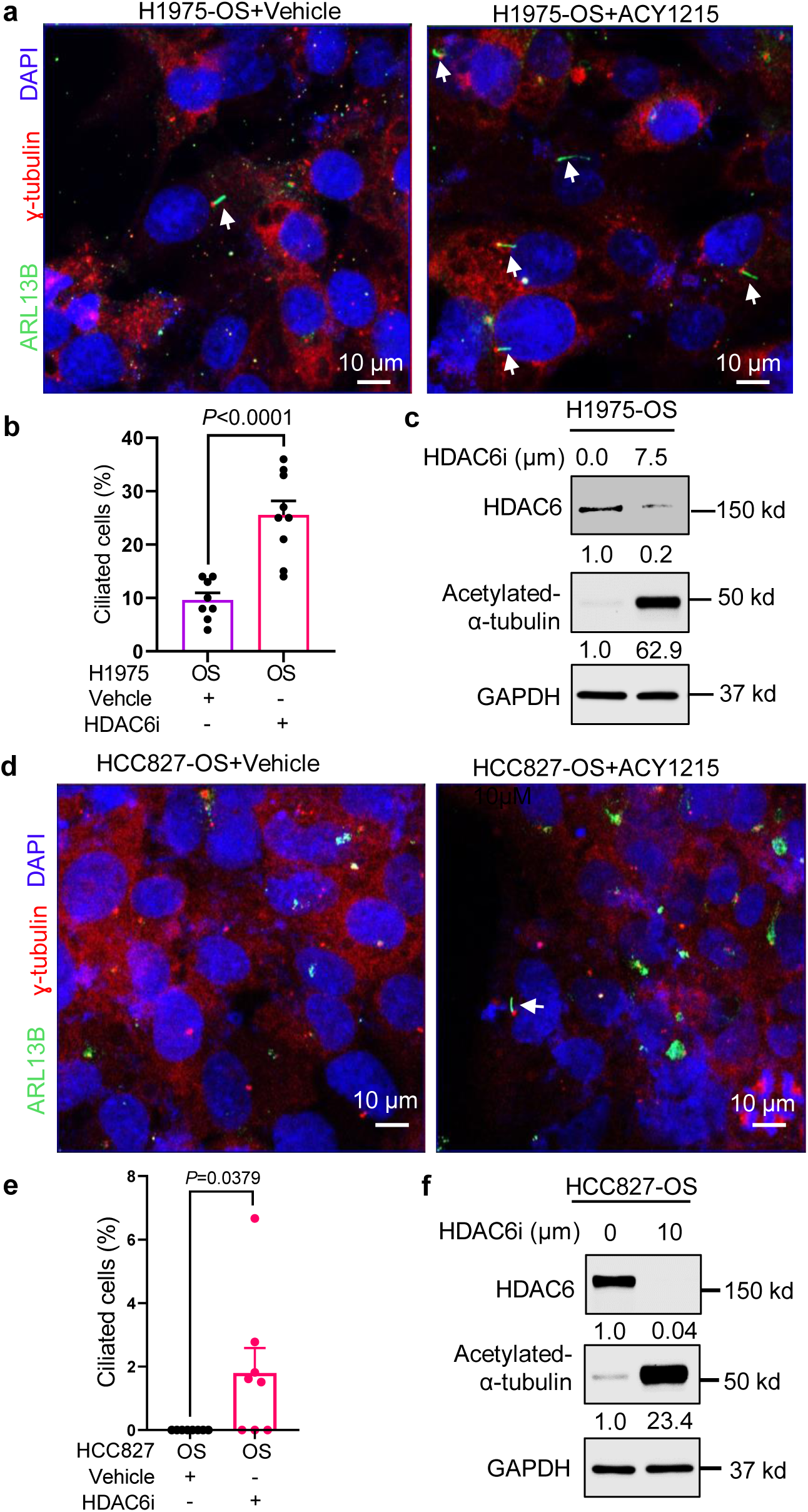
Pharmacologic inhibition of HDAC6 increases primary cilia frequency in osimertinib-sensitive human lung cancer cells. **a-e,** Serum-starved H1975-OS and HCC827-OS were treated with vehicle or the indicated amount of HDAC6 inhibitor ACY1215 (HDAC6i) for 72 h. **a, d,** Cells were fixed & subjected to immunofluorescence staining to detect ARL13B (green) to mark cilia, ɣ-tubulin (red) for centrioles, & DAPI (blue) for nuclei and imaged on a Zeiss LSM900. **b, e,** Quantification of **a** and **d**. One-way ANOVA analysis followed by a Bonferroni test to adjust for multiple comparisons. **c, f,** Cells were lysed & immunoblotted with the indicated antibodies. Densitometry results are shown as numbers below the bands. Images in **a** & **d** are representative of three independent experiments.

**Figure 3.**
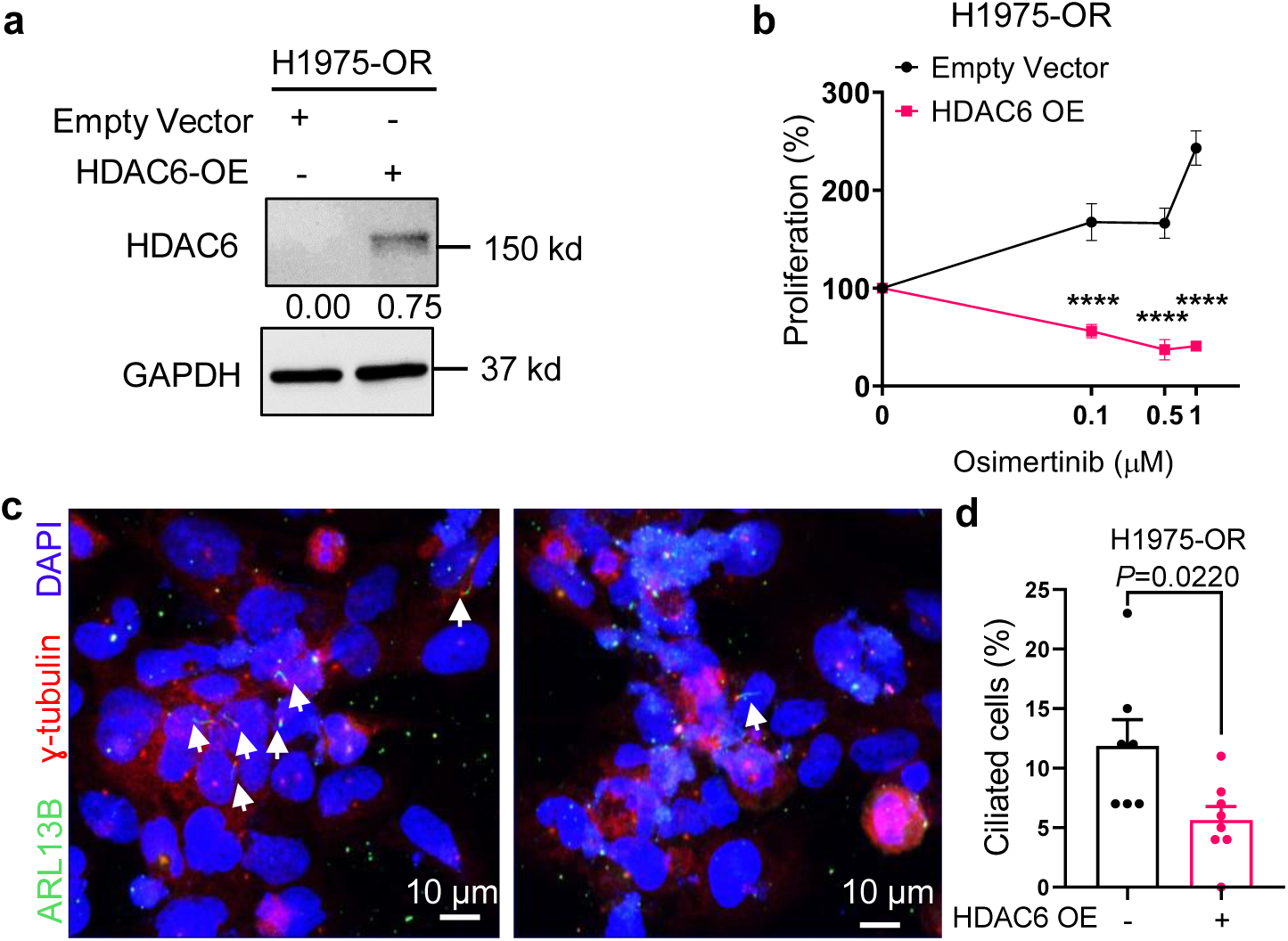
Exogenous HDAC6 overexpression reduces cilia formation & sensitizes osimertinib-resistant NSCLC cells to osimertinib. **a,** Lysates from empty vector– or HDAC6 plasmid–transfected H1975-OR cells were immunoblotted to detect HDAC6 or GAPDH. **b,** CellTiter-Glo® proliferation assays were performed in empty vector– or HDAC6 vector–transfected H1975-OR cells 120 h after treating the cells with 0, 0.1, 0.5, or 1.0 μM osimertinib. Two-way ANOVA & Tukey’s test for multiple comparisons. \*\*\*\**P* < 0.0001. **c,** Serum-starved H1975-OR cells were transfected with empty vector or HDAC6 plasmid, fixed, & subjected to immunofluorescence (IF) staining to detect ARL13B to mark cilia & ɣ-tubulin for centrioles. **d,** Quantification of **c.**

### Genetic inhibition of ciliogenesis reverses osimertinib resistance

Given that intraflagellar transport (IFT) proteins are essential for formation and function of primary cilia^38^, we silenced the IFT88 protein to assess whether genetic inhibition of ciliogenesis sensitizes osimertinib-resistant cells to the osimertinib (**Fig. 4a**). As expected, formation of cilia decreased upon shRNA-mediated IFT88 protein knockdown in both H1975-OR and HCC827-OR cells relative to corresponding control cells that had been transduced with control LacZ shRNA (**Fig. 4b-c; Supplementary Fig. 3**). In the presence of osimertinib, IFT88-knockdown H1975-OR cells exhibited significantly reduced proliferation relative to control cells (**Fig. 4d**), suggesting that inhibition of ciliogenesis sensitizes H1975-OR cells to the effects of osimertinib on cellular proliferation. Correspondingly, we observed a decreased IC_50_ in osimertinib-treated H1975-OR cells with IFT88 knockdown (0.8092 μm) relative to similarly treated cells with LacZ knockdown (2.546 μm; **Fig. 4e-f**).

**Figure 4.**
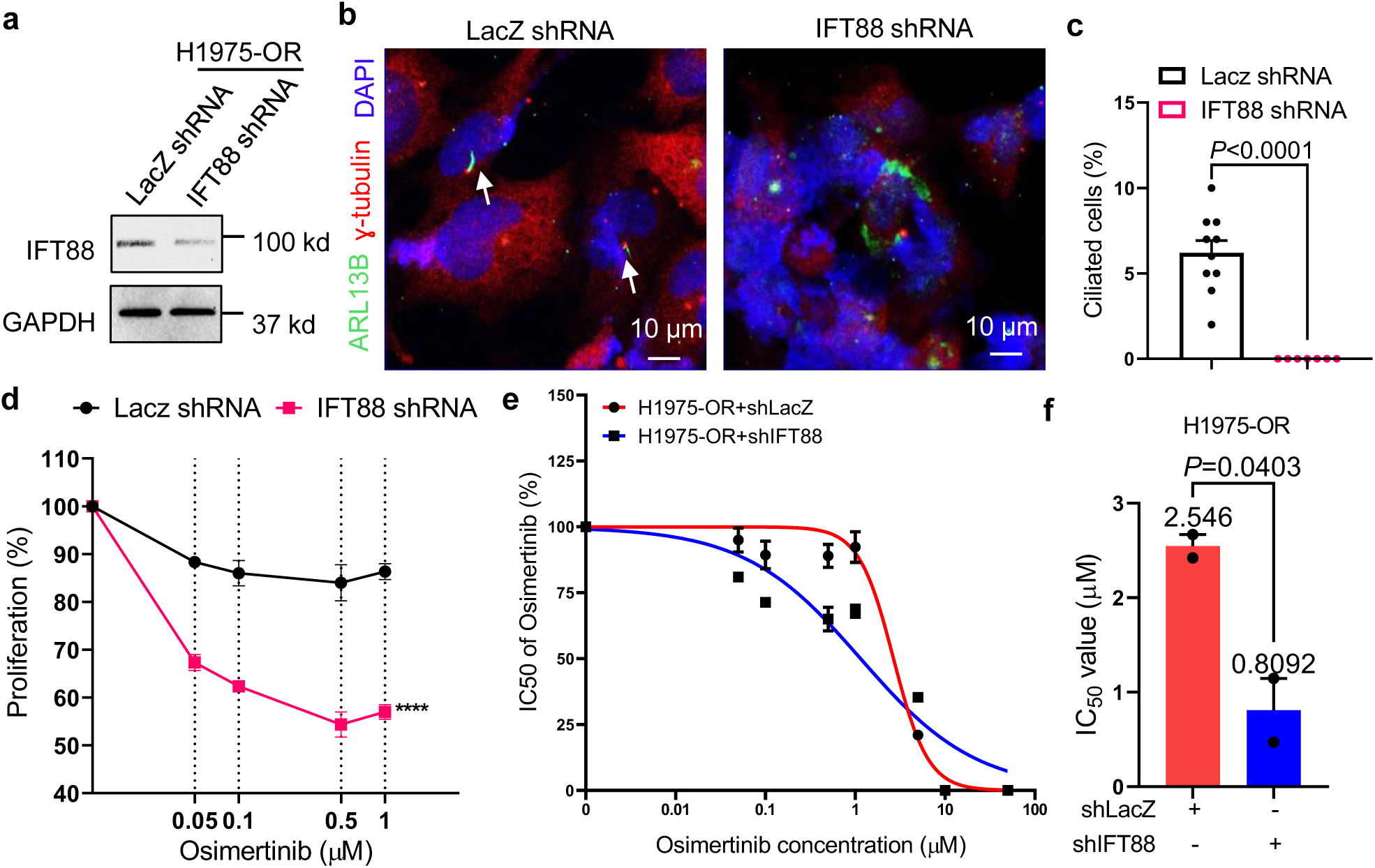
Genetic inhibition of ciliogenesis by knocking down IFT88 protein sensitizes osimertinib-resistant cells to EGFR TKI–mediated inhibition of proliferation. **a,** H1975-OR cells stably transduced with lentivirus encoding LacZ- or IFT88-specific shRNA were lysed & immunoblotted. **b,** Immunofluorescence of fixed serum-starved H1975-OR cells was performed to detect ARL13B (green) to mark cilia, ɣ-tubulin (red) for centrioles, and DAPI (blue) for nuclei using a Zeiss LSM900 confocal microscope. White arrows denote cilia. **c,** Quantification of **b**. Unpaired t-test. **d,** CellTiter-Glo® proliferation assays were performed in H1975-OR cells stably transduced with lentivirus encoding LacZ- or IFT88-specific shRNA 120 hours after treating the cells with 0, 0.05, 0.1, 0.5, or 1.0 μM osimertinib. Two-way ANOVA & Tukey’s test for multiple comparisons. \*\*\*\**P* < 0.0001. **e,** IC50 of osimertinib was determined by CellTiter-Glo® proliferation assays in H1975-OR cells. **f,** Comparison of IC50 of osimertinib in H1975-OR cells stably transduced with lentivirus encoding either LacZ-or IFT88-specific shRNA.

To assess whether inhibition of cilia formation reverses osimertinib resistance in vivo, we challenged mice with H1975-OR cells transduced with control LacZ shRNA or IFT88 shRNA lentivirus stably expressing luciferase. Specifically, we injected 1.5 × 10^6^ cells (suspended in PBS with Matrigel) into the left thorax of nonobese severe combined immunodeficient γ (NSG) mice. Once lung tumors were established after 1 week, the mice were administered 5 mg/kg osimertinib in 1% carboxymethyl cellulose solution via oral gavage once every 3 days for 2 weeks. Luciferase imaging was used to assess lung tumor progression (**Fig. 5a; Supplementary Fig. 4**). Tumors from mice harboring IFT88-knockdown H1975-OR xenografts had greater sensitivity to osimertinib, as evidenced by their reduced proliferation, than tumors from mice challenged with control H1975-OR cells, as shown by luciferase imaging (**Fig. 5b**) and Ki-67 staining of histological sections of fixed lung tumor tissue harvested from the mice (**Fig. 5c**). As expected, lung tumors derived from mice challenged with IFT88-knockdown H1975-OR xenografts exhibited less cilia formation than tumors derived from mice challenged with control xenografts (**Fig. 5d, e**). In a complementary approach, NSG mice were subcutaneously implanted with HCC827-OR cells that had been transduced with lentivirus encoding IFT88- or control LacZ-specific shRNA and treated with osimertinib over the course of 8 weeks. We observed substantially reduced tumor growth in mice challenged with IFT88-knockdown lung tumor cells relative to mice that received LacZ-knockdown control cells, as evidenced by decreased tumor volume (**Fig. 6a**), size (**Fig. 6b**), and weight (**Fig. 6c**), as well as reduced cell proliferation (**Fig. 6d-e**). The IFT88-silenced lung tumors had significantly fewer cilia than LacZ-knockdown control tumors (**Fig. 6f-g**). Taken together, these in vivo findings suggest that inhibition of ciliogenesis by IFT88 knockdown sensitizes osimertinib-resistant cells to drug-induced inhibition of proliferation.

**Figure 5.**
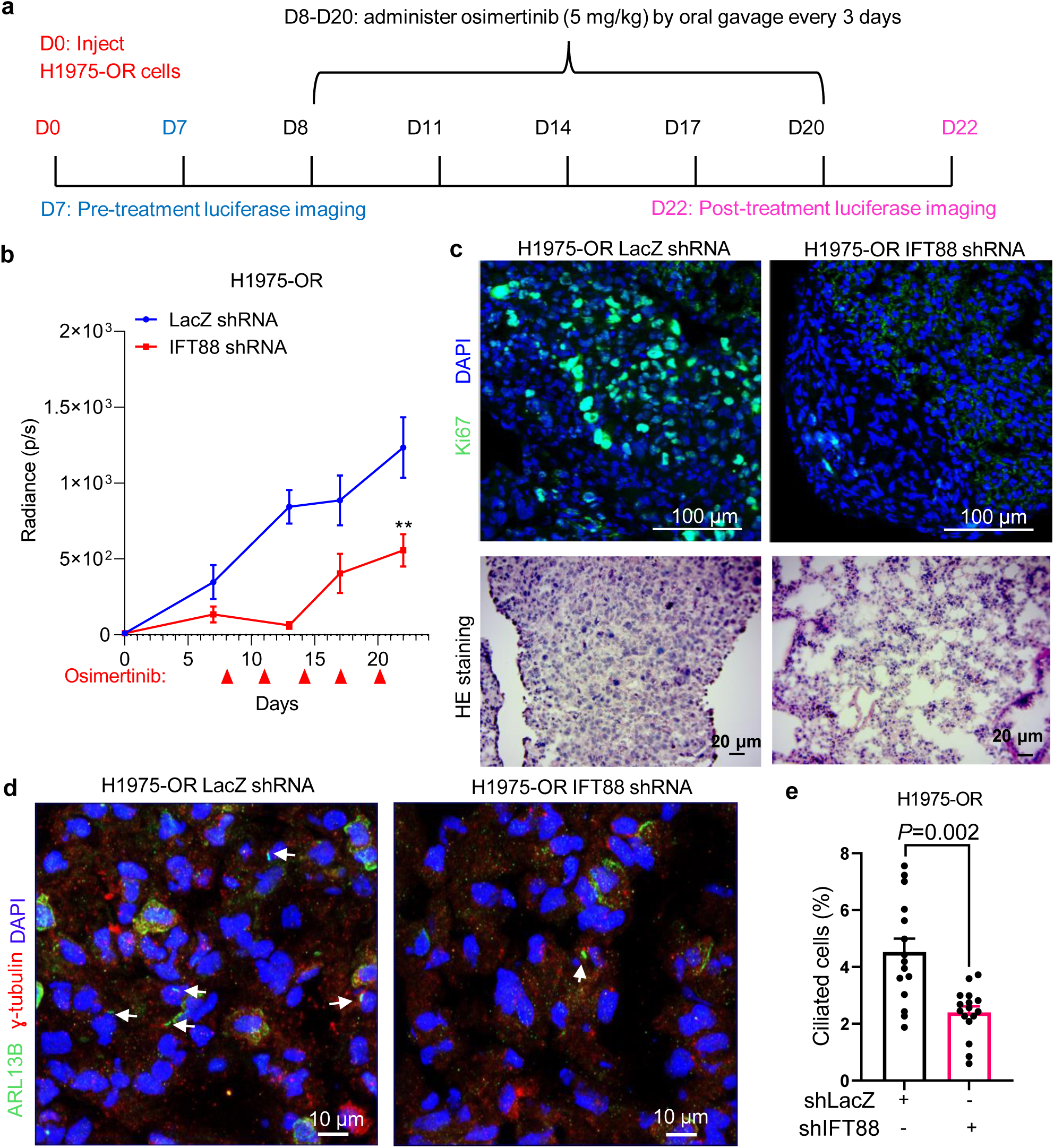
Inhibition of primary cilium formation sensitizes osimertinib-resistant lung tumors to EGFR TKI in mice challenged with orthotopic H1975-OR xenografts. **a,** Luciferase-labeled human H1975-OR cells transduced with LacZ shRNA or IFT88 shRNA were injected into the left thorax of SCID mice. Mice were imaged for luminescence, administered 5 mg/kg osimertinib on the indicated days, and re-imaged near the experimental endpoint. **b,** The growth of the tumor was monitored through luminescence, and the average luminescence intensity at each time point was plotted. Total luminescence intensity (photon count) was calculated using live imaging software. Error bars indicate SEM. \*\**P* < 0.01, two-way ANOVA followed by Tukey’s multiple comparisons test. **c,** (Top) Immunofluorescence was performed using a monoclonal Ki-67 antibody (green) and DAPI (blue) on cryo-sectioned lung tissues obtained from mice (n=3 per group) of the human lung tumor orthotopic model. Scale bar, 100 μm. (Bottom) H&E staining of serial sections of the related tissue. **d,** Immunofluorescence was performed using ARL13B (green) to mark cilia, ɣ-Tubulin (red) for centrioles, and DAPI (blue) for nuclei on cryo-sectioned lung tissues obtained from mice of the human lung tumor orthotopic model. Arrows indicate cilia. Scale bar, 10 μm. **e,** Quantification of **d**. Unpaired t-test.

**Figure 6.**
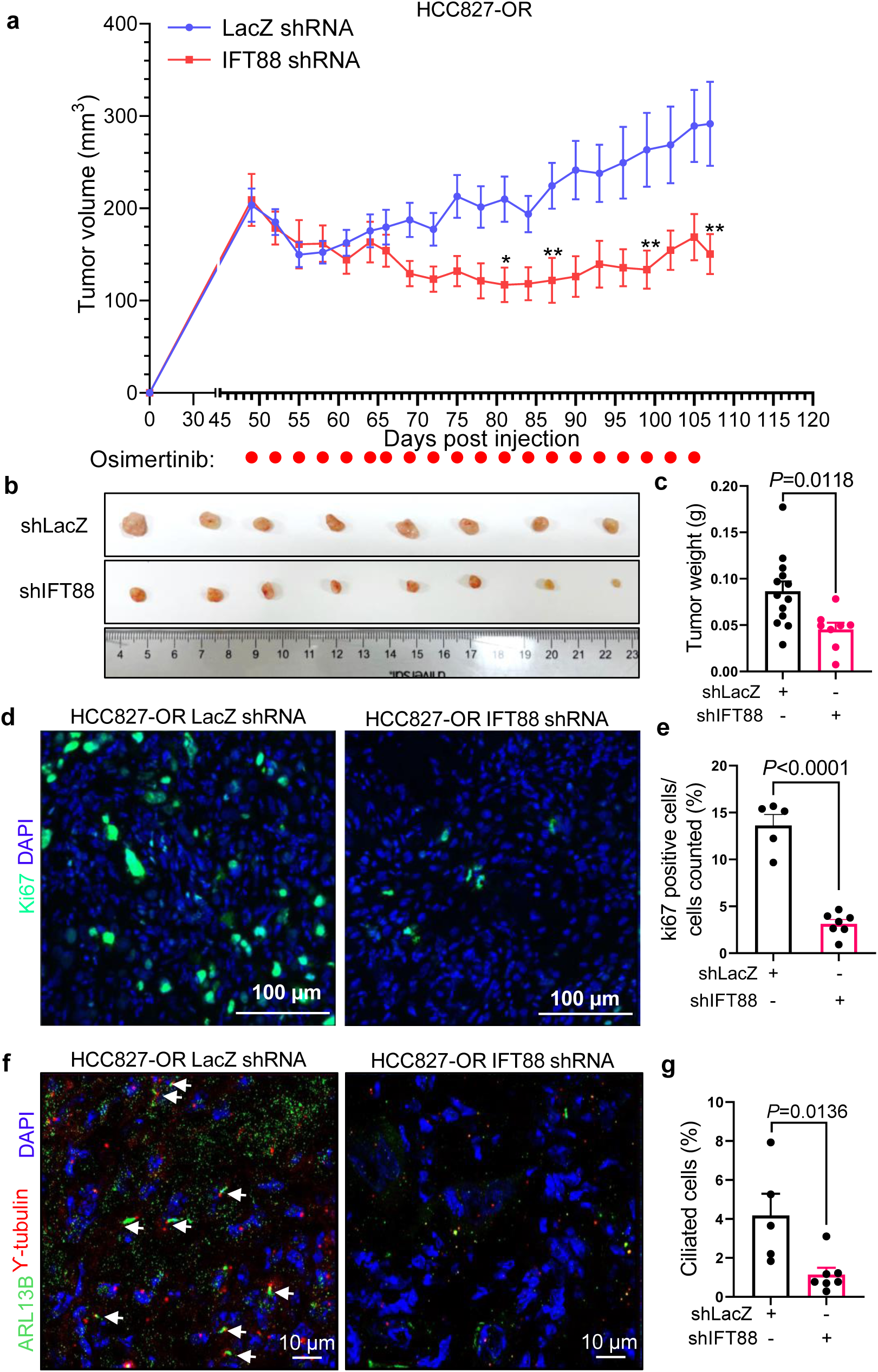
Inhibition of primary cilium formation by silencing IFT88 sensitizes osimertinib-resistant lung tumors to EGFR TKI in mice challenged with subcutaneous HCC827-OR xenografts. **a**, Human HCC827-OR cells transduced with LacZ shRNA or IFT88 shRNA were injected subcutaneously into the left flanks of SCID mice. Mice were administered 5 mg/kg osimertinib on the indicated days. Tumor volume was measured every 5 days with luminescence imaging. **b**, Images of tumors harvested from mice at the conclusion of the experiment. **c**, Tumor weight at the conclusion of the experiment. Unpaired t-test. **d**, Immunohistochemistry was performed using a monoclonal Ki-67 antibody (green) and DAPI (blue) on cryo-sectioned lung tissues (n = 5 mice in the LacZ shRNA group and n = 7 in the IFT88 shRNA group) obtained from subcutaneous human lung tumor model mice. **e**, Quantification of Ki-67 positive cells in **d**. Unpaired t-test. **f**, Immunohistochemistry was performed using ARL13B (green) to mark cilia, ɣ-tubulin (red) for centrioles, and DAPI (blue) for nuclei on cryo-sectioned lung tissues obtained from mice of the human lung tumor model. **g**, Quantification of **f**. Unpaired t-test.

### Decreased intracellular sodium influx reduces ciliogenesis and sensitizes human NSCLC to osimertinib

EGF-induced intracellular sodium influx regulates EGFR trafficking through increased tubulin acetylation^39^. Increased sodium influx through sodium ionophores or by inhibition of Na,K-ATPase mimics the effects of EGF on EGFR trafficking through HDAC6 inactivation and increased acetylation of tubulin^39^. Therefore, we hypothesized that increased sodium influx during osimertinib-induced inhibition of EGFR signaling promotes cilia formation by generating sustained HDAC6 inactivity and greater tubulin acetylation as cancer cells acquire resistance to osimertinib. To test this hypothesis, we induced increased sodium influx in H1975-OS cells with the sodium ionophore gramicidin A (**Supplementary Fig. 5a-b**). We observed a reduction in HDAC6 protein expression (**Supplementary Fig. 5c**), an increase in α-tubulin acetylation (**Supplementary Fig. 5c**), and increased primary cilium formation relative to vehicle controls (**Supplementary Fig. 5d-e**). Given that the ultimate goal is reversing osimertinib-refractory NSCLC progression, we tested whether reduced sodium within the cell inhibits ciliogenesis and sensitizes NSCLC cells to osimertinib treatment. We administered the cAMP agonist dibutyryl-cAMP (DBC)^40^, or vehicle, to H1975-OR cells to reduce intracellular sodium by stimulating the Na,K-ATPase pump. In H1975-OR cells, 100 µM DBC reduced intracellular sodium (**Fig. 7a-b**), increased HDAC6 expression (**Fig. 7c**), decreased α-tubulin acetylation (**Fig. 7c**), decreased ciliogenesis (**Fig. 7d-e**), and sensitized H1975-OR cells to osimertinib-mediated reductions in proliferation (**Fig. 7f**). Likewise, in mice harboring orthotopic H1975-OR tumors, DBC reduced sodium influx (**Fig. 7h-i**) and sensitized the osimertinib-resistant tumors to the drug’s anti-proliferative effects, as evidenced by decreased tumor growth (**Fig. 7g**; **Supplementary Fig. 6**). Collectively, our data suggests that increased sodium influx during the development of osimertinib resistance promotes cilium formation through HDAC6 inactivation and increased acetylation of tubulin.

**Figure 7.**
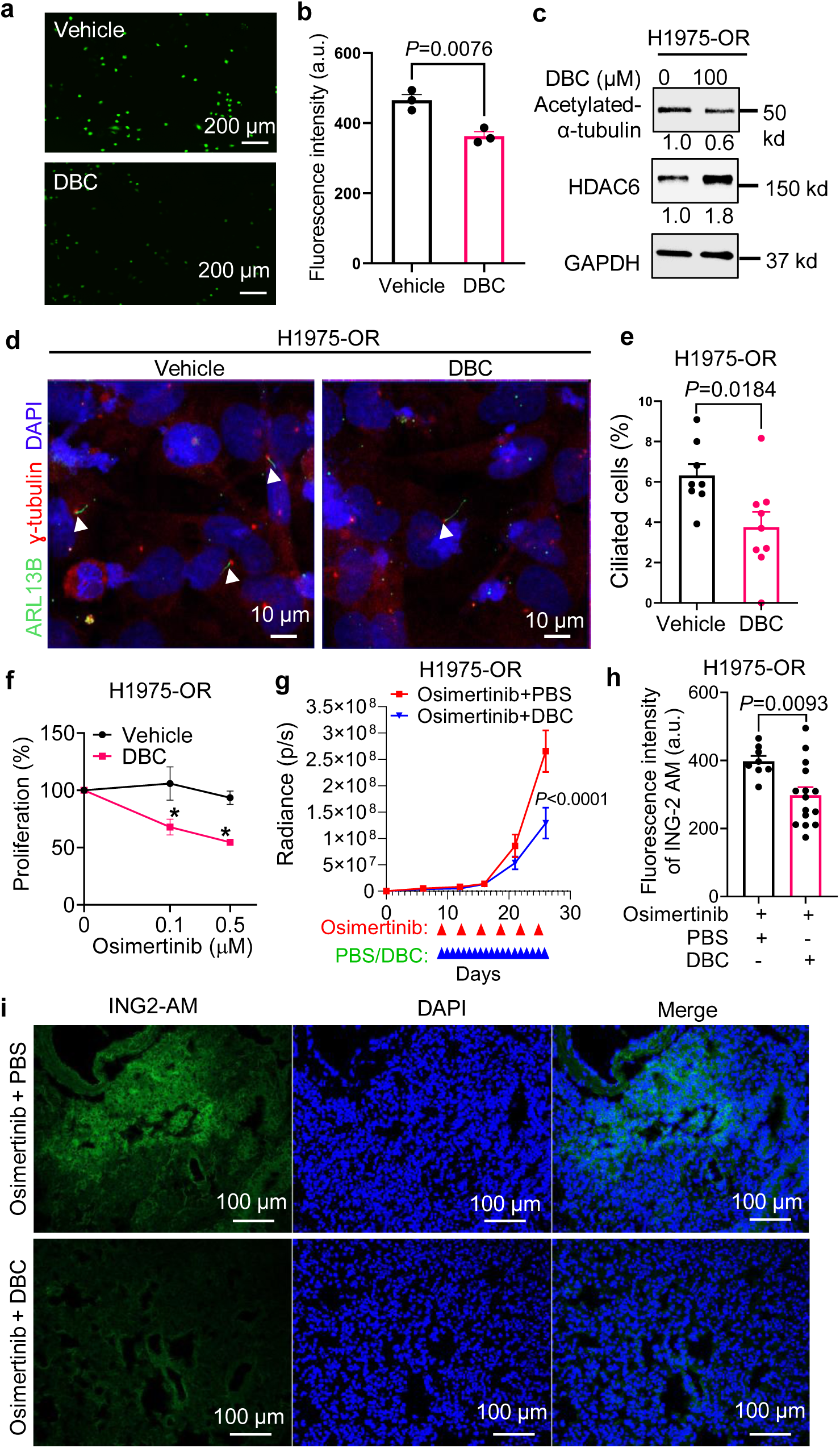
Decreased intracellular sodium influx mediated by DBC reduces ciliogenesis & sensitizes human NSCLC cells to osimertinib. **a,b**, Serum-starved H1975-OR cells were treated with 100 μM dibutyryl-cAMP (DBC), a cAMP agonist that stimulates the Na, K-ATPase pump, resulting in reduced intracellular sodium influx. Fluorescence was quantified with a Bio-Tek synergy Neo2 plate reader. Unpaired t-test was used for analysis. **c**, Serum-starved H1975-OR cells treated with 100 μM DBC for 24 h were lysed for immunoblotting. **d**, Serum-starved H1975-OR cells treated with 100 μM DBC for 24 h were subjected to IF staining to detect ARL13B (green) to mark cilia, ɣ-tubulin (red) for centrioles, and DAPI (blue). **e**, Quantification of **d**; unpaired t test. **f**, CellTiter-Glo® proliferation assays were performed 120 h after osimertinib treatment. Two-way ANOVA & Tukey’s test for multiple comparisons. \**P* < 0.05. **g**, Luciferase-labeled human H1975-OR cells were injected into the left thorax of SCID mice. Mice were orally administered 5 mg/kg osimertinib on the indicated days, and injected intraperitoneally with PBS or DBC every day. Luciferase was imaged at regular intervals to monitor tumor growth. Error bars indicate SEM. Two-way ANOVA followed by Bonferroni’s multiple comparisons test. **h,i,** Histochemistry was performed using the sodium indicator ING-2 AM (green) for sodium influx and DAPI (blue) for nuclei on cryo-sectioned lung tissues obtained from mice of the human lung tumor model. **h** shows quantification of **i**.

## Discussion

Identifying therapeutic strategies to prevent or overcome resistance to first-line osimertinib remains an urgent unmet clinical need. The molecular basis of osimertinib resistance has yet to fully emerge, because first-line osimertinib treatment^7,8^ became standard of care only in 2018 in the US. In our study, we have shown that primary cilium formation is enhanced in NSCLCs resistant to osimertinib. This enhancement is mediated through reduced HDAC6 activity and increased acetylation of α-tubulin. This finding is supported by our observations of increased ciliogenesis upon HDAC6 inhibition and reduced cilia formation upon overexpression of HDAC6. IFT88 is required for the formation and functionality of primary cilia^38^. Our data shows that genetic ablation of IFT88 decreases primary cilia and reduces osimertinib-refractory lung cancer progression both in vitro and in vivo. Mechanistically, our findings in mice demonstrate that inhibition of intracellular sodium influx through drug-induced stimulation of the Na, K-ATPase pump results in decreased cilia formation, increased sensitivity to osimertinib, and reduced tumor progression, suggesting that intracellular sodium influx positively regulates ciliogenesis and osimertinib resistance in human EGFR-mutated NSCLC cells.

HDAC6 regulates primary cilia disassembly and shortening by destabilizing microtubules through deacetylation of α-tubulin in the ciliary axoneme^33^. In zebrafish, BBIP10 localizes to the primary cilium and physically interacts with Hdac6^41^. Loss of the deacetylation enzyme Hdac6 through mutation promotes acetylation of cytoplasmic tubulin^42^, and inhibition of Hdac6 restores microtubule acetylation in BBIP10-depleted cells. A study in embryonic human stem cells and induced pluripotent cells reported that ceramide prevents activation of HDAC6 by cytosolic αPKC and Aurora kinase A, which promotes acetylation of tubulin in primary cilia in stem cells and neural progenitors and, potentially, neural processes^43^. In cancer cells, depletion of METTL3-mediated m(6)A modification leads to abnormally elongated cilia via suppressing HDAC6-dependent deacetylation of axonemal α-tubulin, ultimately attenuating cell growth and cervical cancer development^44^. In cholangiocarcinoma, microRNA dysregulations lead to HDAC6 overexpression and ciliary loss, while the use of HDAC6 inhibitors restored ciliary expression^33,45^. Taken together, these results show that it is well established that reduced HDAC6 activity and increased acetylation of α-tubulin is a hallmark of primary cilia formation. Indeed, here we show that HDAC6 inactivity, elevated α-tubulin acetylation, and increased ciliogenesis is associated with acquired osimertinib resistance in NSCLC.

Given that over 60% of NSCLCs express EGFR with gain-of-function or activating mutations leading to constitutive tyrosine kinase activity, finding the mechanism is important to support the development of novel anticancer agents that target EGFR^46^. Inhibition of HDAC6 has been shown to reduce tumor growth in models of glioblastoma^47^ and cholangiocarcinoma^33^. Based on these prior observations, we speculated that NSCLC cells that develop resistance to the EGFR TKI osimertinib would exhibit decreased HDAC6 activity, increased acetylation of α-tubulin, and increased primary cilia formation, a hypothesis that was supported by our current results.

IFT is a highly conserved process common to all ciliated or flagellated eukaryotic cells, in which membrane-bound vesicular cargo is carried along the microtubular axoneme in association with non–membrane-bound protein complexes^48^. In cisplatin-induced acute kidney injury, a decrease in IFT88 expression has been linked to impaired, shortened primary cilia^49^. Correspondingly, genetic inhibition of IFT88 through siRNA has been shown reduce the length of cilia^49^. Several mouse models have also established that IFT88 is required for primary cilia formation. Ift88-null mutations in mice (*Ift88^tm1Rpw^*, *Ift88^tm^*^1^*^.1Bky^*, and *Ift88^fxo^*) are embryonic lethal about when organogenesis begins^50^, but The Oak Ridge Polycystic Kidney (ORPK) mouse relies on insertional mutagenesis to partially disrupt the expression and function of IFT88, resulting in a hypomorphic allele (designated *Ift88^Tg737Rpw^*) that enables these homozygous mutant mice to survive into adulthood in the absence of functional primary cilia^51^, making them a good model for studying the pathogenesis of human ciliopathies. Tissue-specific deletion of *Ift88* in retinal pigment epithelium in another mouse model of primary cilia loss^52^, suggests that defective cilia promote local wound healing. In our studies reported here, we impaired cilium formation in human osimertinib-resistant H1975 and HCC827 cells through stable shRNA-mediated knockdown of IFT88; orthotopically xenografted the luciferase-labeled, IFT88-knockdown cells into SCID mice; administered osimertinib; and observed that loss of primary cilia sensitizes osimertinib-resistant lung tumors to drug-induced cytotoxicity, suggesting that primary cilium formation contributes to EGFR TKI resistance in vivo. However, we cannot discard the ciliary independent functions of IFT88^53–55^. Nevertheless, our findings are supported by prior work showing increased cilia in HCC4006 NSCLC cells that are resistant to the first-generation EGFR TKI erlotinib and A549 lung cancer cells that are resistant to chemotherapy^20^.

EGF-induced intracellular sodium influx regulates EGFR trafficking through increased tubulin acetylation^39^. Increased sodium influx mimics the effects of EGF on EGFR trafficking by inactivating HDAC6 and increasing acetylation of tubulin^39^. Given that we observed increased primary cilia formation, reduced HDAC6 activity, and increased acetylation of α-tubulin in EGFR-mutated, osimertinib-resistant NSCLC cells, we proposed that increased sodium influx during inhibition of osimertinib-induced EGFR signaling promotes cilia formation by sustaining HDAC6 inactivity, leading to greater tubulin acetylation as cancer cells acquire resistance to osimertinib. This hypothesis was supported by our results that a drug-induced reduction in intracellular sodium influx increased HDAC6 activity, decreased α-tubulin acetylation, decreased ciliogenesis, and sensitized H1975-OR cells to osimertinib-mediated reductions in proliferation in vitro and in vivo (**Fig. 7**).

Pharmacological inhibition of intracellular sodium influx may therefore represent a promising approach to inhibiting ciliogenesis and, consequently, to sensitizing EGFR TKI–refractory NSCLCs to osimertinib. This approach is appealing because currently no specific small-molecule inhibitor of primary cilia formation has been reported, although some small molecules have been shown to inhibit cilium assembly while simultaneously impacting other cellular functions. Given that the activation of Hedgehog signaling requires functional cilia, and changes in ciliogenesis regulate Hedgehog pathway activation, it is not surprising that studies seeking to identify Hedgehog pathway inhibitors discovered repressors of primary ciliogenesis. The small molecule HPI-4 (eventually renamed Ciliobrevin A) represses ciliogenesis by inhibiting Dynein-2-dependent retrograde IFT; however, HPI-4 also blocks the ATPase activity of dynein in the cytoplasm, which alters a wide range of cilium-independent cellular functions, including mitotic spindle formation^56,57^. Two other Hedgehog signaling inhibitors, CA1 and CA2, repress primary cilia formation in a direct fashion through disruption of IFT^58^, but likely affect dynein-dependent cellular processes in the cytoplasm and microtubule cytoskeletal organization^59^. Several protein kinases have been found to be important regulators of primary cilium formation. Small-molecule inhibitors of fibroblast growth factor receptor 1 (FGFR1) repress ciliogenesis^20,60^ and sensitize erlotinib-resistant HCC4006 human lung cancer cells to this first-generation EGFR TKI^20^. Unfortunately, FGFR1 inhibitors (i.e., SU5402 and BGJ398) exhibit promiscuous off-target effects on other FGF receptors and protein kinases from different families, leading to widespread cilium-independent regulation of cellular processes^59^. Cyclin-dependent kinases also regulate primary ciliogenesis. CDK5 potently promotes primary cilia formation, and two CDK5 inhibitory molecules, roscovitine and its analog (S)-CR8, repress ciliogenesis directly through inhibition of IFT and indirectly through disruption of the microtubule cytoskeleton. Inhibition of some other molecular targets by drugs has unexpectedly been shown to also repress ciliogenesis, including HSP90 (ganetespib), neddylation (MLN4924), and tyrosine kinases (sunitinib, erlotinib), but again these inhibitors affect cellular processes beyond regulation of ciliogenesis. Although these inhibitors of ciliogenesis represent invaluable tools for basic research studies designed to better understand the role of primary cilia formation in development and pathological conditions, including cancer, the clinical goal is to deliver drugs only to specific tissues where primary cilia contribute to disease pathology and progression to minimize the side effects of the drug, particularly in healthy tissues. Tumor-restricted delivery of pharmacological inhibitors of intracellular sodium influx or selectively cytotoxic agents that restrict intracellular sodium loading could be viable strategies to suppress ciliogenesis, thereby restoring osimertinib sensitivity in EGFR TKI-resistant NSCLC cells.

In conclusion, our study suggests that increased primary cilia formation on the surface of lung tumor cells promotes osimertinib-refractory NSCLC progression. Our in vitro and in vivo data show that targeted inhibition of ciliogenesis can prevent or overcome osimertinib resistance, which may lead to breakthroughs in preventing progression of EGFR TKI–refractory lung adenocarcinoma and may lead to future clinical trials to treat NSCLC.

## Methods

### Cell lines and culture

Osimertinib-sensitive H1975 parental (H1975-OS) and H1975 osimertinib-resistant (H1975-OR)^32^ human NSCLC cell lines were a generous gift from Dr. Anthony C. Faber at Virginia Commonwealth University. HCC827-OR were generated by incubating HCC827-OS in serial increases in concentration of osimertinib and expanding single colonies. H1975-OS and HCC827-OS were grown in RPMI-1640 complete medium (Corning) containing 10% fetal bovine serum (FBS; Sigma, Cat no. F0926), 1% penicillin/streptomycin antibiotics (Corning, Cat no. 30-002-CI), and 25 μg/mL plasmocin prophylactic (Invitrogen, Cat no. NC9886956). RPMI-1640 complete medium with 1 μM osimertinib (Selleckchem, Cat no. S7297) was used to grow H1975-OR and HCC827-OR. HEK-293T cells were purchased from ATCC and maintained in DMEM (Corning) containing 10% FBS; Sigma), 1% penicillin/streptomycin antibiotics (Corning), and 25 μg/mL plasmocin prophylactic (Invitrogen).

### Immunofluorescence studies

Cells were seeded at the density of 1 × 10^5^ per well on EMD Millipore Millicell EZ Slides (Sigma Millipore) pretreated with collagen I (Corning, Cat no. 354231) and grown in RPMI-1640 complete medium. The cells were starved in medium without FBS for 36 hours, when they achieved full confluency, then fixed in 4% paraformaldehyde (Boston BioProducts,) and stained for cilia markers ARL13B (Proteintech, Cat no. 17711-1-AP, 1:200) and ɣ-tubulin (Sigma, Cat no. T5326, 1:200) or acetylated-α-tubulin (Sigma, Cat no. T6793, 1:200). Nuclei were stained with 4’, 6-diamino-2-phenylindole (DAPI; Cell Signaling Technology, Cat no. 8961). Alexa Fluor™ 488 goat anti-rabbit IgG (Invitrogen, A-11008) and Alexa Fluor™ 594 goat anti-mouse IgG (Invitrogen, A-11005) were used. All images were captured on a Zeiss LSM900 confocal microscope using a 63×/1.4 NA objective and ciliated cells were analyzed using ImageJ software (https://imagej.net/software/fiji/).

Frozen sections of the lung tissues harvested from the in vivo orthotopic lung cancer model were fixed in 4% paraformaldehyde) and stained for cilia markers ARL13B (Proteintech, 1:200), ɣ-Tubulin (Sigma, 1:200), and proliferation marker Ki67 (CST, Cat no. 12202S, 1:200) 24 hours after sectioning. Nuclei were stained with DAPI.

### Immunoblotting

Cells were lysed in RIPA buffer (Millipore) containing protease inhibitors (Roche) and phosphatase inhibitors (Sigma-Aldrich). Proteins were separated via 4-20% gradient SDS-PAGE (Bio-Rad), transferred to membranes, and blocked with 5% bovine serum albumin (Sigma-Aldrich). After the blocking, the membranes were immunoblotted with one of these primary antibodies: HDAC6 (CST, Cat no. 7558, 1:1000), acetylated-α-Tubulin (Sigma, Cat no. T6793, 1:1000), GAPDH (CST, Cat no. 5174, 1:1000), or tubulin (Santa Cruz Biotechnology, sc-5286, 1:500). Following overnight incubation in blocking buffer (TBST containing 5% bovine serum albumin), the blots were washed in TBST and incubated with secondary antibodies as described previously^61^. Antibody-reactive protein bands were detected by enzyme-linked chemiluminescence (Thermo Fisher Scientific) using a ChemiDoc MP Imaging System (Bio-Rad) and quantified using ImageJ software (https://imagej.net/software/fiji/) to obtain densitometry values.

### Proliferation assay

Cells were counted by using a DeNovix CellDrop FL Cell Counter and seeded onto 96-well plates at a concentration of 30k per well. Proliferation assays were conducted with a CellTiter-Glo® 2.0 Cell Viability Assay (Promega, Cat no. G9242) 5 days after osimertinib treatment. Data were analyzed as the ratio of luminescence values of osimertinib-treated to -untreated cells.

### Transient transfection

The previously described purified HDAC6-Flag plasmid^62^ was quantified using Nanodrop 2000 and transiently transfected following the instructions for the Lipofectamine™ 3000 Transfection Reagent (L3000008). The effect of overexpression was verified by immunoblotting against HDAC6.

### Generation of stable cell lines

To prepare the lentivirus, LacZ shRNA and IFT88 shRNA cloned into pLKO.1 plasmids (Sigma, TRCN0000145198) were transfected into 293T cells along with their corresponding packaging plasmids. The lentivirus was isolated from cell culture medium 48 h after transfection and used immediately to transduce H1975-OR or HCC827-OR cells. Stable IFT88 knockdown cells were used for experiments following confirmation of IFT88 protein silencing by immunoblotting.

### Chemical reagents

Gramicidin A from *Bacillus brevis* (Cat no. 50845) and dibutyryl-cAMP (DBC, Cat no.16980-89-5) were purchased from Sigma. H1975-OS cells were starved, and gramicidin A was added at 200 nM when they achieved 80% confluency. H1975-OR cells were starved and treated with 100 μM DBC. Following 48 hours of treatment, both cell lines were subjected to immunoblotting, immunofluorescence, and proliferation assays.

### In vivo orthotopic lung cancer model

The 6- to 8-week-old pathogen-free NSG mice were maintained under specific pathogen-free conditions in a temperature-controlled room with a 12-h light/12-h dark cycle. The 8- to 12-week-old male and female mice were orthotopically injected with 1.5 × 10^6^ luciferase-labeled human H1975-OR lung cancer cells stably transduced with LacZ or IFT88 shRNAs suspended in 50 μl PBS and 30 μl Matrigel. After lung tumors were established, as seen in the luminescence images captured using an In-Vivo Xtreme xenogen imaging system (Bruker), the mice were administered 5 mg/kg osimertinib every 3 days. Tumor growth was tracked by luminescence imaging every 5 days. At the end of the experiment, mice were euthanized by CO_2_ inhalation, and the lungs were perfused with PBS and harvested. A portion of the lung was preserved in O.C.T. (Scigen, #4586K1) immersed in 2-methylbutane (Sigma, #32631) that had been pre-cooled in dry ice. Cryo-sections were collected using a cryostat (Leica). Bruker molecular imaging software (Living imaging 4.7.3) was used to calculate luminescence intensity (total flux [photons/second]) in each mouse. All animal studies were performed in accordance with the protocols approved by the University of Minnesota Institutional Animal Care and Use Committee.

For the mice treated with PBS or DBC, male and female NSG mice at 8 to12 weeks old were orthotopically injected with 1.5 × 10^6^ luciferase-labeled human H1975-OR lung cancer cells. After lung tumors were established, as seen on in vivo luminescence images, the mice were administered 5 mg/kg osimertinib every 3 days by oral gavage and PBS/DBC every day by intraperitoneal injection.

### In vivo subcutaneous lung tumor model

Eight- to twelve-week-old male or female NSG mice were subcutaneously injected with 2 x 10^6^ HCC827-OR cells stably transduced with LacZ shRNA or IFT88 shRNA suspended in 100 μl PBS and high-concentration Matrigel (Corning, #354248). To monitor the growth of the tumors, tumors were measured using electronic digital Venier calipers every 3 days, and tumor volume was calculated using the formula (length x width^2^)/2. Mice were treated with 5 mg/kg osimertinib when the tumor volume was around 200 mm^3^. At the end of the experiment, tumors were extirpated, photographed, weighed, and preserved in O.C.T. for immunofluorescence analysis.

### Hematoxylin and eosin (H&E) staining

Frozen sections of the lung tissues harvested from mice of the in vivo orthotopic lung cancer model were fixed in 4% paraformaldehyde for 30 minutes and washed with deionized water before the nuclei were stained with IHC-Tek Mayer’s Hematoxylin Solution (IHC World, Cat no. IW-1400). The slides were washed in water, blued in 0.5% ammonia solution for 30 seconds, and immersed in 95% ethanol briefly, after which the cytoplasm was stained with eosin Y disodium salt (mpBio, Cat no. 195700). After being washed with water, immersed in 95% ethanol for 1 minute, dipped 10 times in 100% ethanol, and immersed in xylene twice for 2 min each, the slides were mounted with SignalStain® Mounting Medium (CST, Cat no.14177S) using Corning® glass coverslips (Sigma, Cat no. CLS2975225). All images were captured on a Zeiss Apotome 2 microscope using a 20×/0.8 NA Plan Apochromat objective and processed using ImageJ software.

### Intracellular sodium measurement

H1975-OS or H1975-OR cells were seeded in a Greiner 96-well plate at a concentration of 4,000 cells/well. Twenty-four hours later, cells were starved in RPMI-1640 basal medium overnight followed by the addition of 200 nM gramicidin A or 100 μΜ DBC. At the indicated time point, the medium was removed and 100 μl of 2 μg/μl ING-2 AM (Ion Bioscience, Cat. no. 2011G) diluted in 1X Pluronic F-127 solution (Ion Bioscience, Cat. no. 7601A) was added to the cells, which were then incubated for 60 minutes in a cell culture incubator.

Fluorescence images were acquired by using an Incucyte® S3 equipped with a green filter and 10× objective. The intensity of the fluorescence was read with a plate reader (Bio-Tek synergy Neo2; excitation/emission: 525 nm/545 nm).

### Statistical analysis

An unpaired, two-tailed Student’s t-test or one-way ANOVA was used to compare differences between groups as indicated, and values of *P* < 0.05 were considered significant. The Bonferroni or Tukey test was used to correct for multiple comparisons when necessary. Data is expressed as mean ± SEM and is representative of multiple independent experiments.

## Acknowledgements

We thank The Hormel Institute and its staff for administrative, shared equipment, animal facility, and institutional support. We are grateful to Dr. Kishor Pant, Dr. Estanislao Peixoto, and Seth Richard in Dr. Sergio Gradilone’s Lab for sharing reagents. We thank Dr. Naomi Ruff for providing editorial support.

## Author Contributions

L.W. conducted most of the in vitro cell line-based experiments, including immunofluorescence studies, generation of stable cell lines, immunoblotting, and proliferation assays. S.K.A. and R.S. contributed to the immunofluorescence experiments. L.W., A.P., S.K.A., and R.S. captured and analysed immunofluorescence images. S.K.A. generated osimertinib-resistant HCC827-OR cells. L.W., A.P., S.K.A., and L.H.H. managed the mouse colony. L.W. and L.H.H. performed tumour studies in mice, conducted murine in vivo imaging, tumour measurements, and necropsy. L.W., A.P., S.K.A., S.A.G., and L.H.H. provided technical and scientific support. L.W. and L.H.H. performed experimental troubleshooting, reviewed relevant scientific literature, critically analyzed data, prepared figures, and wrote the manuscript. L.H.H. conceived the aims, led the project, and acquired funding to complete the reported research.

## Funding

This work was supported by a Research Scholar Grant RSG-21-034-01-TBG from the American Cancer Society, an LCRF Research Grant on Oncogene Driven Lung Cancer (Award ID: 992504) from the Lung Cancer Research Foundation and EGFR Resisters, and a Paint the Town Pink Award sponsored by The Hormel Institute Internal Grants Program to L.H.H. This work was also supported by The Hormel Foundation to L.H.H. and S.A.G. and by the National Institutes of Health R01DK13281 grant funding to S.A.G.

## Supplementary figure legends

**Supplementary Figure 1.**
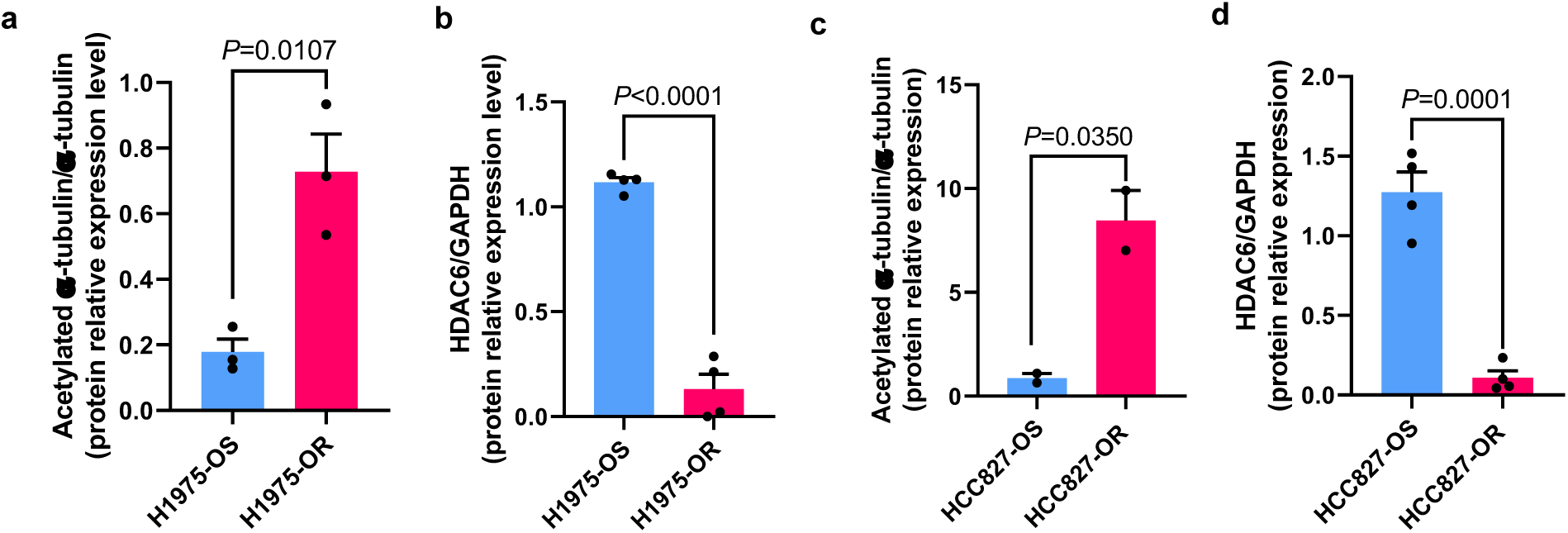
Densitometry analysis of immunoblots that appear in Figure 1. **a-d,** Serum-starved osimertinib-sensitive (H1975-OS, HCC827-OS) and osimertinib-resistant (H1975-OR, HCC827-OR) cells were lysed and immunoblotted with the indicated antibodies. Densitometry was performed on the immunoblotting images by quantifying acetylated α-tubulin relative to α-tubulin, and HDAC6 relative to GAPDH, as indicated. Reported values in the bar graph represent the mean of three biological replicates (i.e., each black closed circle represents a densitometry value from one biological replicate). Unpaired t-test.

**Supplementary Figure 2.**
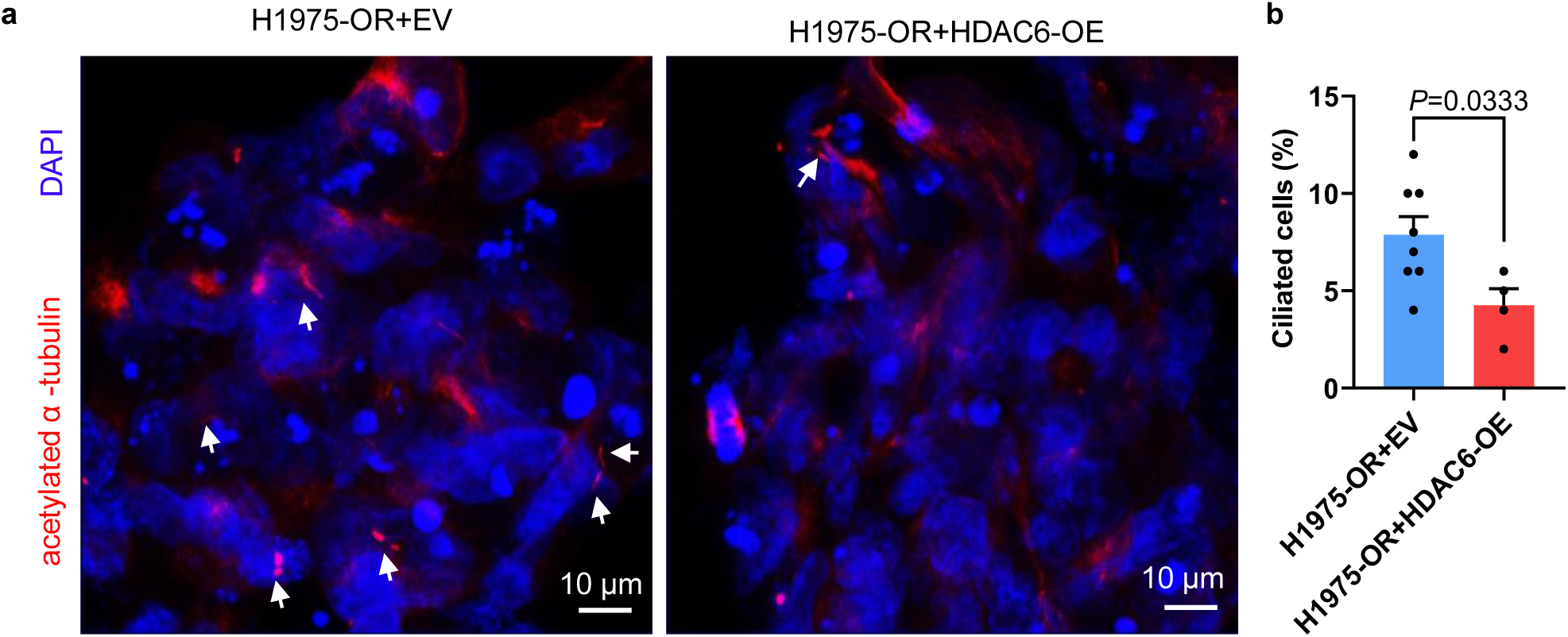
Exogenous HDAC6 overexpression reduces cilia formation. **a,** Serum-starved H1975-OR cells were transfected with empty vector or HDAC6 plasmid, fixed, & subjected to immunofluorescent staining to detect acetylated-α-tubulin to mark cilia. **b,** The quantification **a**. Unpaired t-test.

**Supplementary Figure 3.**
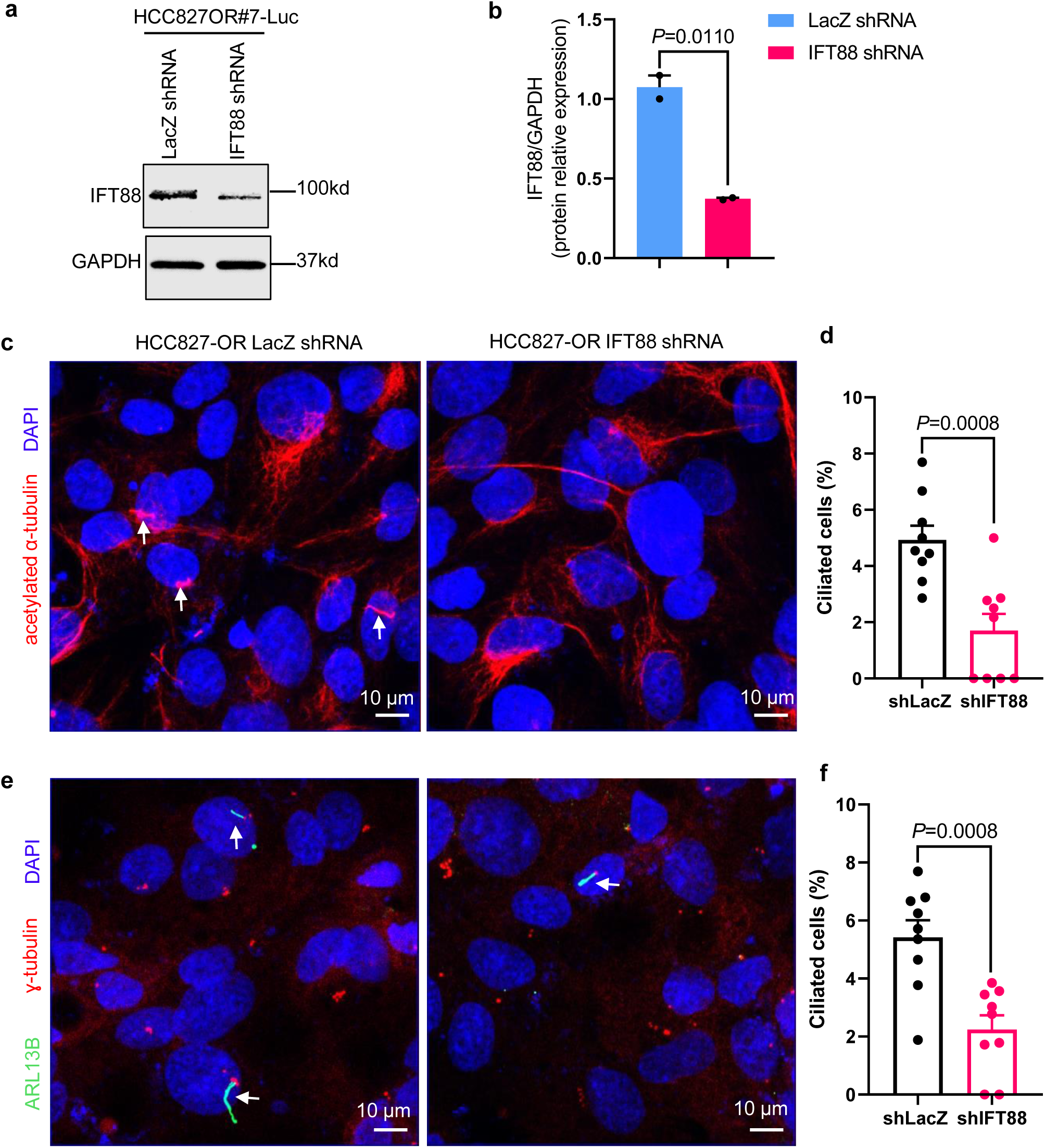
Genetic inhibition of IFT88 protein reduces ciliogenesis in osimertinib-resistant NSCLC cells. **a**, HCC827-OR cells stably transduced with lentivirus encoding IFT88- or LacZ-specific shRNA were lysed & immunoblotted. **b**, Quantification of blots. **c**, Immunofluorescent staining of fixed, serum-starved HCC827-OR cells to detect acetylated-α-tubulin (red) to mark cilia and DAPI (blue) for nuclei. White arrows denote cilia. **d**, Quantification of **c**. Unpaired t-test. **e**, Immunofluorescence of fixed serum-starved HCC827-OR cells was performed to detect ARL13B (green) to mark cilia, ɣ-tubulin (red) for centrioles, and DAPI (blue) for nuclei using a Zeiss LSM900 confocal microscope at 63×. White arrows denote cilia. **f**, Quantification of **e**. Unpaired t-test.

**Supplementary Figure 4.**
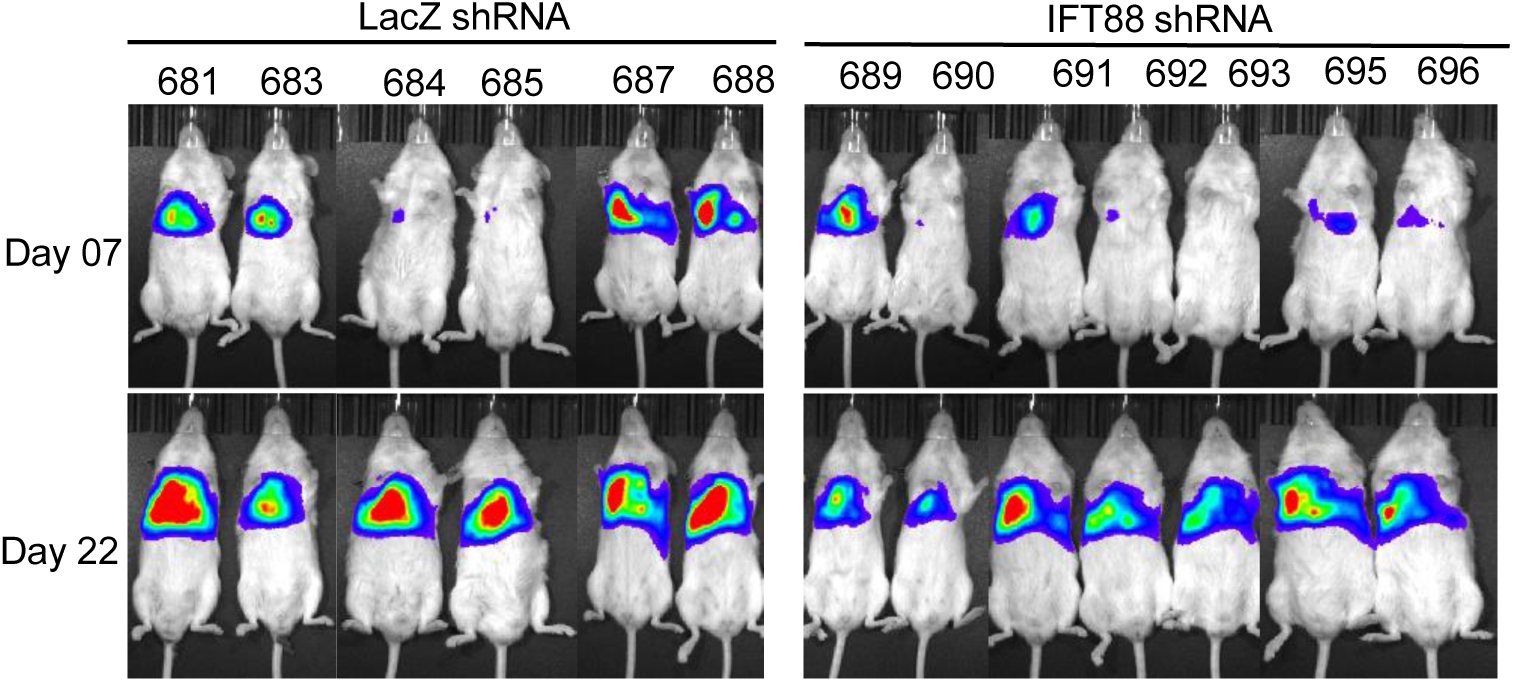
Genetic inhibition of ciliogenesis by silencing IFT88 protein sensitizes osimertinib-resistant cells to EGFR TKI–mediated inhibition of proliferation. Luciferase-labeled human H1975-OR cells were injected into the left thorax of SCID mice. Mice were xenogen imaged, and representative luminescence images are depicted showing lung tumors immediately prior to osimertinib treatment at day 7 and following osimertinib treatment at the conclusion of the study at day 22.

**Supplementary Figure 5.**
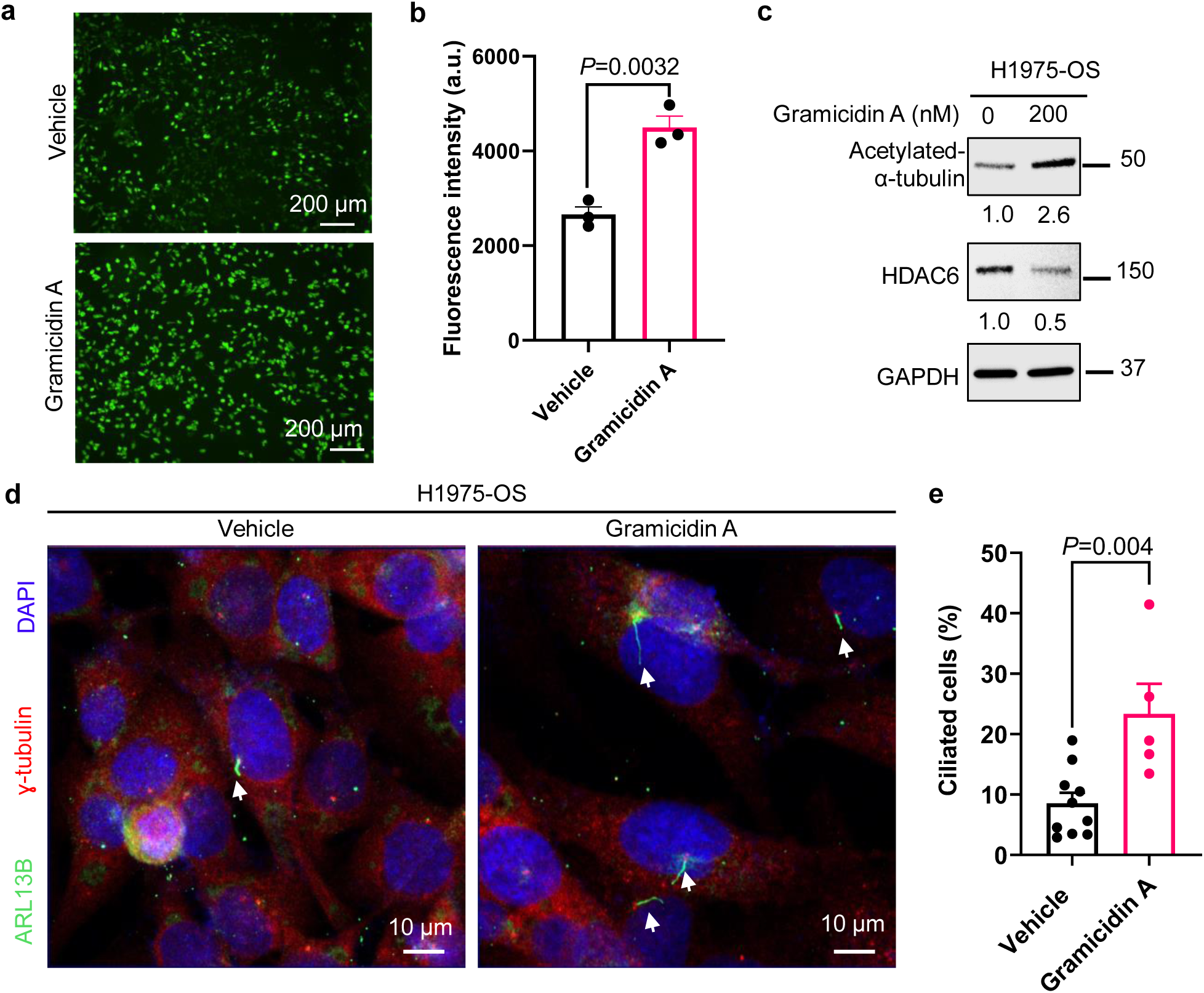
Intracellular sodium influx regulates ciliogenesis in human EGFR-mutated NSCLC cells. Serum-starved H1975-OS cells were treated with 200 nM gramicidin A for 48 h. **a,b**, Gramicidin treatment resulted in increased intracellular sodium influx, as detected with 2 μg/μl ING-2 AM. An unpaired Student t-test was used for analysis. **c**) The cells were lysed for immunoblotting. **d**) The cells were subjected to IF staining to detect ARL13B (green) to mark cilia, ɣ-tubulin (red) for centrioles, and DAPI (blue). **e**) Quantification of **d**; unpaired t test.

**Supplementary Figure 6.**
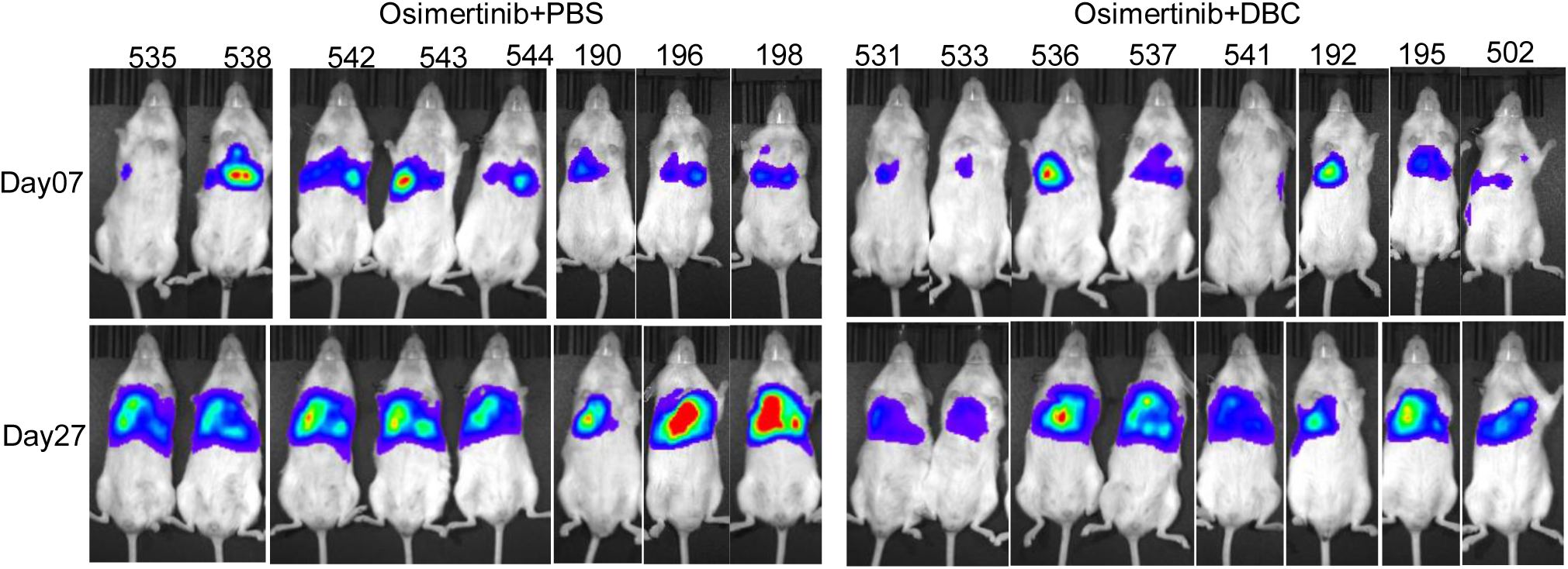
Decreased intracellular sodium influx mediated by DBC reduces ciliogenesis and sensitizes human NSCLC cells to osimertinib. Luciferase-labeled human H1975-OR cells were injected into the left thorax of SCID mice. Mice were imaged for luminescence. Representative images are shown from day 7, right before treatment, and day 27, close to the end point of the experiment.

## Notes

### Competing Interest Statement

The authors have declared no competing interest.

